# Use of orthogonal split inteins to direct surface expression of specific metabotropic glutamate receptor hererodimer pairs

**DOI:** 10.64898/2026.11.23.768648

**Authors:** Viviana Marolda, Tyler Couch, Tyler W. McCullock, Amanda Newell, John D. Lueck, Paul J. Kammermeier

## Abstract

Metabotropic glutamate receptors (mGluRs) are class C G protein coupled receptors that function as obligate, covalently linked homo- and heterodimers in some combinations. The pharmacological responses of heterodimers to selective ligands are difficult to predict, and due to the tendency to homodimerize, heterodimers cannot be easily expressed in isolation. Here, we developed a split-intein-based approach to preferentially direct the plasma membrane expression of defined mGluR heterodimers, enabling their pharmacological characterization with minimal homodimer interference. Our strategy employs orthogonal split inteins flanking an endoplasmic reticulum retention/internalization sequence within the receptor C-terminal tail. Association of receptors carrying complementary intein modules promotes protein trans-splicing, removing the trafficking restriction and allowing receptor delivery to the cell surface. Following a systematic screening, the orthogonal Avi PRP8 and Mja KlbA inteins (designated C and F) were selected to generate complementary constructs. Co-expression of these constructs produced functional responses from multiple group II and III mGluR combinations, whereas individually expressed receptors generally lacked detectable signaling. Pharmacological characterization of mGlu2/4 using subtype-selective agonists supported the preferential expression of functional heterodimers over homodimers. Furthermore, glutamate dose-response profiles of most CF/FC receptor combinations closely resembled those obtained using the heterodimer-selective CODA-RET assay, supporting the ability of this approach to distinguish heterodimer-mediated signaling. Independent HiBiT-based surface-expression measurements confirmed partner-dependent membrane delivery while revealing receptor-specific limitations in retention efficiency, particularly for mGlu8-containing constructs. Together, these findings establish orthogonal split inteins as a tool for the selective functional and pharmacological characterization of mGluR heterodimers while minimizing interference from homodimer signaling.

## Introduction

Metabotropic glutamate (mGlu) receptors are class C G protein coupled receptors with widespread expression in the central nervous system^1,2^. These receptors are important regulators of synaptic efficacy and neural excitability, and many have been implicated as therapeutic targets for pathologies such as autism, schizophrenia, and Parkinson’s disease^3–7^. There are 8 mammalian mGlu genes that function as obligate dimers, either as homodimers or as heterodimers in certain combinations^1,8^. Interestingly, pharmacological responses to both allosteric and orthosteric compounds are dramatically altered in different mGlu heterodimers, but isolating heterodimer signaling is complicated and has generally involved at least some alterations of the receptors, particularly in the C-terminal ends, leading to the possibility that receptor function or response to ligands may not represent the true effects on more fully wild type receptors. Further, many of these approaches are BRET based and are not amenable to examination of signaling to downstream effectors exclusively from each heterodimer, since they do not prevent homodimer formation.

To address this problem, a relatively effective strategy has been employed to express strict heterodimer expression including generation of mGluR-GABA_B_ receptor chimeras in which the C-tails of GABA_B_Rs replace a portion of the mGluR tail to force heterodimer expression through masking of internal ER retention sequences through coiled-coil interactions^9,10^. This approach has some utility but requires a robust alteration of the native C-terminal sequence, which could plausibly affect receptor signaling and potentially pharmacology^11^. Alternatively, a split luciferase strategy called complimented donor-acceptor resonance energy transfer (CODA-RET)^12^ has also been employed with good effect in which the fragments of luciferase are fused to corresponding mGluR C-tails yielding enhanced BRET signals when co-expressed with a fluorescent acceptor-tagged G protein^13,14^. We have recently used a similar approach using the low affinity Nanoluciferase fragments LgBit and SmBit paired with a miniG_o_-venus to take a similar approach to examine heterodimer specific G protein recruitment to all of the likely group II (mGlu2 and 3) and III (mGlu4, 7, and 8) mGluR heterodimer pairs^15^. However, this approach also has some limitations. First, like the GABA_B_R tail chimer approach, it requires significant mutagenesis of the C-tail of each receptor, and because the method isolates the BRET signal to heterodimers but does not prevent homodimer expression, it cannot be employed to examine heterodimer specific downstream signaling effects.

To overcome these limitations, we have engineered a new approach to heterodimer composition control in which C-tail ER retention and internalization sequences are flanked by orthogonal split intein^16–18^ sequences such that upon dimerization, splicing between the two inteins removes the retention/internalization sequences and simultaneously transfers the native C-tail of the partner receptor to its N-terminal domain, leaving only a few additional amino acids as a ‘scar’ to the otherwise nearly wild type heterodimer. Here we describe the process of identifying suitable intein pairs, tagging of mGluRs, optimization and testing before demonstrating that this system can be used to generate nearly all potential group II and III mGluR heterodimer pairs.

## Materials and Methods

### Cell culture and maintenence

HEK293T cells were maintained in DMEM supplemented with 10% fetal bovine serum (FBS), 1% of penicillin/streptomycin (GIBCO) and 1% of GlutaMax (GIBCO) at 37°C in a humidified 5% CO2 atmosphere. To harvest cells from the plate for subculturing, they were washed with PBS 1X and incubated with trypsin/EDTA for 5 minutes at 37°C. Afterwards, cells were resuspended in fresh media, and the appropriate passage of cells was performed, usually ranging between 1:5 to 1:10. Sequences of the ER retention/internalization module as well as all of the N-and C-inteins are shown in Appendix A.

### G-Protein Dissociation NanoBRET Assay

G-protein activation was monitored in live cells using a NanoBRET-based assay that detects the activation-dependent dissociation of Gβγ from GαoA and its subsequent recruitment to the membrane-tethered C-terminal fragment of GRK3 fused to NanoLuc (masGRK3ct-Nluc), as described ^19–21^. Gβγ was labeled using a split-Venus system, in which complementary fragments of the Venus fluorescent protein were fused to Gβ and Gγ, respectively. HEK293T cells were seeded in 6-well plates at a density of 1.7–2 × 10⁶ cells/well in complete DMEM supplemented with 10% fetal bovine serum (FBS), 1% penicillin-streptomycin (Pen-Strep), and 1% GlutaMAX. After 4 h, the culture medium was replaced with DMEM containing 2% FBS, and cells were transfected with plasmids encoding EAAT3 (400 ng), GαoA (600 ng), Gβ-Venus (200 ng), Gγ-Venus (200 ng), and masGRK3ct-Nluc (50 ng), together with a total of 800 ng of the indicated CF/FC receptor constructs (400 ng of each construct), with or without an N-terminal HiBiT tag. The total amount of transfected DNA was balanced with empty pcDNA3.1 vector when necessary, and polyethyleneimine (PEI) was used as the transfection reagent. Cells were then incubated at 37°C in 5% CO₂ for 24 h. On the day of the experiment, cells were washed with 1× PBS and incubated with 500 µL of 1X PBS containing 5 mM EDTA for 5 min at 37°C in the absence of CO₂. Following incubation, cells were washed three times with Calcium Imaging Buffer (CIB), consisting of 136 mM NaCl, 560 μM MgCl2, 4.7 mM KCl, 1 mM Na2HPO4, 1.2 mM CaCl2, 10 mM HEPES, 5.5 mM Glucose, and then transferred to 1.5-mL microcentrifuge tubes, and centrifuged at 500g for 5 min. Cell pellets were resuspended in 350 µL of CIB, and 25 µL of cell suspension was transferred to each well of a white 96-well microplate (Corning). Cells were then incubated at room temperature for 1 h. Immediately before luminescence acquisition, the NanoLuc substrate Hikarazine-103 (1:125 dilution in CIB; Synthelis Biotech, La Tronche, France) was added to each well, and baseline BRET signals were recorded for approximately 30 s. Serial dilutions of L-Glutamate prepared in CIB were then manually added to the wells. BRET measurements were acquired using a FLUOstar Omega plate reader (BMG Labtech, Ortenberg, Germany) equipped with a dual-emission filter set for NanoLuc donor emission (460–480 nm) and Venus acceptor emission (520–540 nm). The BRET ratio was calculated as the intensity of Venus acceptor emission divided by NanoLuc donor emission (535 nm/475 nm). Ligand-induced changes in BRET (ΔBRET) were calculated by subtracting the baseline BRET ratio recorded before agonist addition from the post-stimulation BRET ratio. Concentration-response curves were generated using absolute ΔBRET values without normalization to the maximal glutamate response and fitted using a three-parameter nonlinear regression model in GraphPad Prism (Boston, MA) to determine glutamate potency (EC₅₀) and maximal response (Emax).

### HiBiT-Based Cell Surface Expression Assay

To assess the cell surface expression of HiBiT-tagged mGlu2CFx constructs, HEK293T cells were seeded in 6-well plates at a density of 2 × 10⁶ cells/well in complete DMEM supplemented with 10% fetal bovine serum (FBS), 1% penicillin-streptomycin (Pen-Strep), and 1% GlutaMAX. After 4 h, the culture medium was replaced with DMEM containing 2% FBS without GlutaMAX, and cells were transfected with the indicated receptor constructs using polyethyleneimine (PEI) as the transfection reagent. For co-transfection experiments, 400 ng of each receptor construct was used, whereas for single-construct transfections, 400 ng of receptor plasmid DNA was supplemented with 400 ng of empty pcDNA3.1 vector to maintain a constant total DNA amount of 800 ng per well. Cells were then incubated at 37°C in 5% CO₂ for 24 h. On the day of the experiment, cells were washed with 1X PBS and incubated with 500 µL of 1X PBS containing 5 mM EDTA for 5 min at 37°C in the absence of CO₂. Following incubation, cells were washed three times with Calcium Imaging Buffer (CIB), consisting of 136 mM NaCl, 560 μM MgCl2, 4.7 mM KCl, 1 mM Na2HPO4, 1.2 mM CaCl2, 10 mM HEPES, 5.5 mM Glucose, and then transferred to 1.5-mL microcentrifuge tubes, and centrifuged at 500 g for 5 min. Cell pellets were resuspended in 300 µL of CIB, and 50 µL of cell suspension was transferred to each well of a white 96-well microplate (Corning). Cells were then incubated at room temperature for 1 h. Approximately 5 min before the start of luminescence recording (t = 0), 50 µL of CIB containing purified Large BiT (LgBiT) protein (1:200 dilution) and Hikarazine 103 (1:125 dilution) was added to each well. At t = 10 min, 11 µL of digitonin solution (4 mg/mL in CIB) was manually added to each well to permeabilize the cells and allow LgBiT access to intracellular HiBiT tags. Luminescence was continuously monitored for an additional 40 min using a FLUOstar Omega plate reader (BMG Labtech). For data analysis, the maximum luminescence recorded before digitonin permeabilization was used as the surface expression signal, whereas the maximum luminescence recorded after permeabilization was used as the total expression signal. Untransfected cells were included in each experiment to determine background luminescence, which was subtracted prior to data analysis. The relative cell surface expression was calculated as the ratio of surface to total luminescence. Background-corrected total relative luminescence units (RLU) were also used as a measure of overall receptor expression. Data are presented as mean ± SEM from at least three independent experiments. Statistical significance was assessed using one-way ANOVA followed by Dunnett’s multiple comparisons test.

## Results

### Intein orthogonality screen

As an initial screen for orthogonality, several split intein pairs were chosen^22^ and subcloned into one of two soluble reporter constructs. Each N-intein was fused to the C-terminal end of monomeric NeonGreen (mNG) with an N-terminal HA tag, and each C-intein was fused to the N-terminal end of nanoluciferase (Nluc) (Fig. 1a). Each combination was expressed in HEK293 cell pairwise. Cell lysates were collected from a set of transfections using the Aov C intein and several N-inteins. These were run on an immunoblot labeled with anti-HA (to detect the mNG-N-intein fragment; green in Fig. 1b) and with anti-NLuc (to detect the C-intein fragment; red in Fig. 1b). Note that the intein splice product, a ∼49 kd predicted fusion protein of HA-mNG and NLuc (upper arrow), was labeled with both antibodies. The blot in Fig. 1b (upper) shows several N-intein constructs (as indicated) in green, paired with the AovC intein product in red (∼37 kd predicted, lower arrow), with the splice product evident in some but not all lanes, indicating that the AovC intein has some limited orthogonality. Note that in lanes with a prominent splice product, the N-intein band is difficult to detect or absent. This is likely due to stoichiometry of expression of the inteins. We also observe a smaller band in each lane where a splice product is evident only labeled by the anti-NLuc antibody. We assume this is a degradation product of the spliced protein in which the HA epitope and perhaps some of the mNG protein is lost. Further examination of the effects of varying the N:C intein ratio is described in Supplemental Fig. 1A.

**Figure 1.**
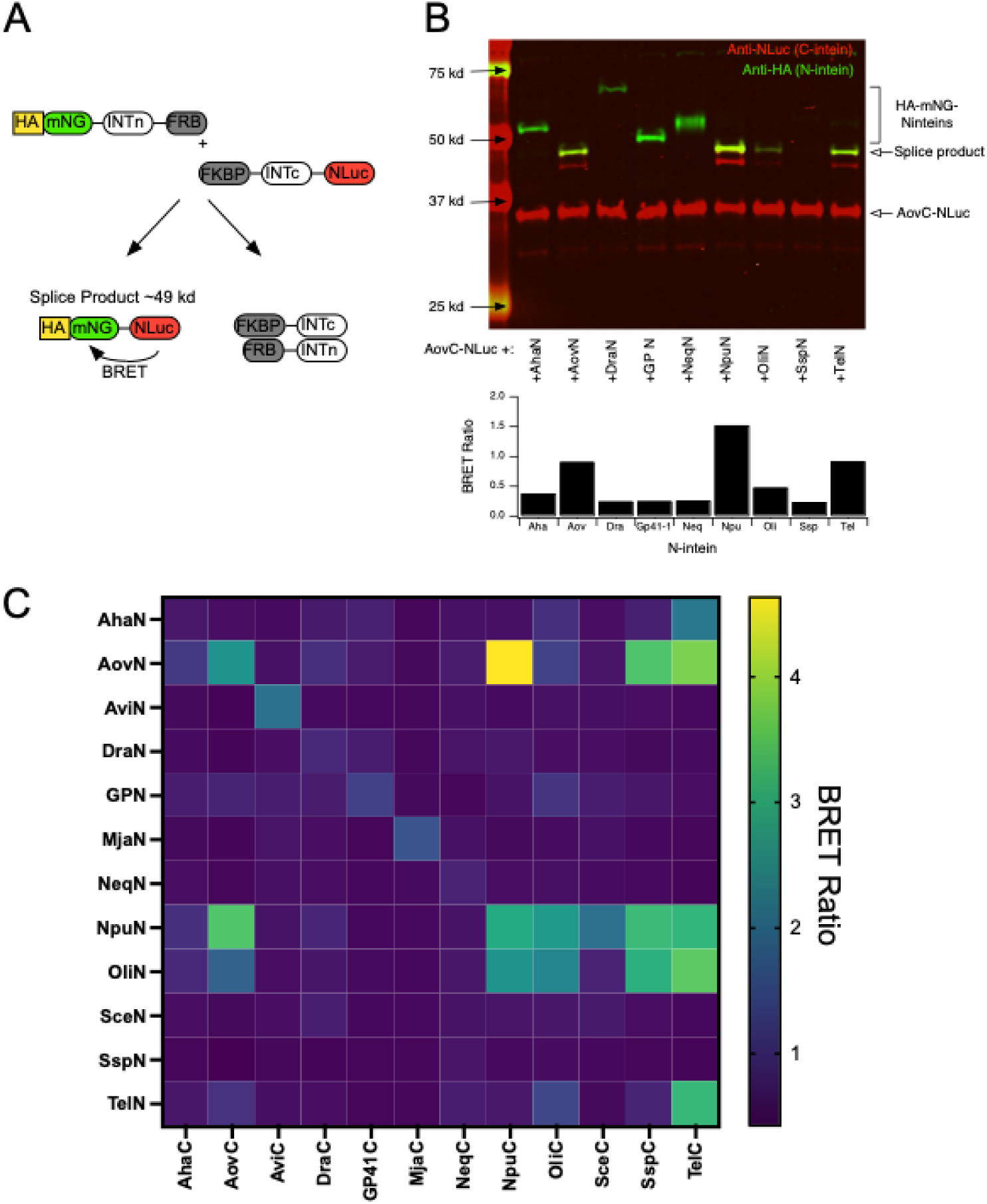
Soluble intein BRET sensors reveal orthagonality. A, Schematic of intein BRET sensors. Each N-intein was fused to an N-extein comprised of an N-terminal HA tag, and monomeric NeonGreen (mNG), and C-terminally to the rapamycin binding domain FRB. Each C-intein was fused to a C-extein comprised of Nanoluciferase (NLuc), and N-terminally to the rapamycin binding domain FKBP12. Following intein association and splicing, an HA-mNG-NLuc fusion protein resulted, leading to an increased BRET signal due to the close proximity of NLuc and mNG. B, Western blot (*upper*) and BRET signal (*lower*) from HEK293T cells transfected with on C-intein sensor (Aovc-Nluc) and several N-intein sensors (tagged with HA-mNG), as indicated. The blot shows anti-HA (green) and anti-NLuc (red). The HA-mNG-NLuc splice product is labeled by both antibodies. Note that the appearance of the splice product corresponds to elevated BRET signals below. C, pooled BRET values from expression of several N-and C-intein sensors.

Next, cells were reserved from each transfection condition and and plated in a 96-well plate and NLuc luminescence as well as mNG luminescence were collected (Fig. 1b, *lower*). The BRET ratios for each combination were collected. BRET values for each sample are shown in Fig. 1b. In general, elevated BRET ratios corresponded well to the appearance of a splice product band in the western blot above.

The BRET sensors were designed with the rapamycin binding domains FRB on the C-terminal end of the N-intein constructs and FKBP12 (“FKBP”) on the N-terminal end of the C-intein constructs. This was intended as a means to facilitate the interaction between each pair of constructs by applying rapamycin. However, rapamycin application had no effect on splicing as assessed by western blot, or on the BRET signal for all but two of the intein pairs, suggesting that for most the association of the N-and C-inteins was a strong enough driver of splicing without rapamycin. The two exceptions were the Neq intein (also called NeqPoln), which exhibited a small increase in BRET between Neqc and some N-inteins, but no evidence of splicing with any (Supplemental Fig. 1B). This result was anticipated since the Neq intein arises from a thermophilic bacteria and requires temperatures above 50 C to splice^23,24^. In addition, the Sce intein (often called VMA) also exhibited rapamycin dependence, as previously reported^25^. Together these data suggest that our BRET sensors can detect intein splicing as an increase in BRET signal, and this increase generally predicts splicing.

Next, a set of 12 N-and C-inteins were chosen for a more complete screen and expressed in every combination with 3-4 replicates (Fig. 1C). Average BRET ratio from these experiments are shown as a heat map. BRET ratio baseline values were around 0.5-0.6, indicating no detectable BRET other than crowding effects. Most N-inteins exhibited elevated BRET (above ∼0.8) when paired with at least one C-intein, with the exception of Neq (discussed above) and Dra. Note that an increased BRET ratio should be representative of splicing, but the upper limit of this value will tend to be lower for pairs in which the C-intein/NLuc construct was stoichiometrically overexpressed compared with the N-intein/mNG construct. In general, most of the inteins tested showed some degree of orthogonality with two (Avi and Mja) showing absolute orthogonality (only splicing with their own partner) with respect to this set of inteins.

### Construction and testing of the intein-ER retention module

Our strategy for engineering an mGluR dimer composition control system was to insert a C-terminal ER retention/internalization sequence into each receptor, flanked by orthogonal split inteins that theoretically should only splice out when two receptors with compatible intein pairs dimerize (Fig. 2A). However, each intein requires a few amino acids in the flanking extein sequence for efficient splicing, which will leave a small ‘scar’ of a few amino acids. ER retention sequences are more effective if located near the plasma membrane, so optimally, the location of the intein/retention modules should be located closely after the 7^th^ transmembrane domain. However, because the C-tail of mGluRs can participate in G protein coupling, we tested a few locations for disruption of signaling by cloning in the Npu intein ‘scar’ into 5 different positions in the mGlu1 C-tail. Position 1.0 was immediately after TM7. Each subsequent position represents an additional 5 AAs. Each construct was tested for function in a plate reader using a NanoBRET system with G_o_a coexpressed (Supp. Fig. 2A and B). With the exception of the scar inserted at position 1.0 (immediately after the last amino acid of TM7), which did not respond to glutamate, receptors with Npu scars inserted at each of the other positions appeared to respond. Dose response curves from three positions (1.1, 1.2, and 1.3) showed that this scar did not detectably alter the pharmacology of the receptor at these positions. Thus, we moved forward using position 1.1 for all further mutants since we did not see disruption of function, but the location remained close to the plasma membrane to optimize ER retention. We also examined scars from three different inteins (*right*) at position P1.1 to evaluate the functional consequences (Supp Fig. 2C). While the Mja scar appeared to enhance efficacy in this experiment (3 replicates), none detectably altered glutamate potency.

**Figure 2.**
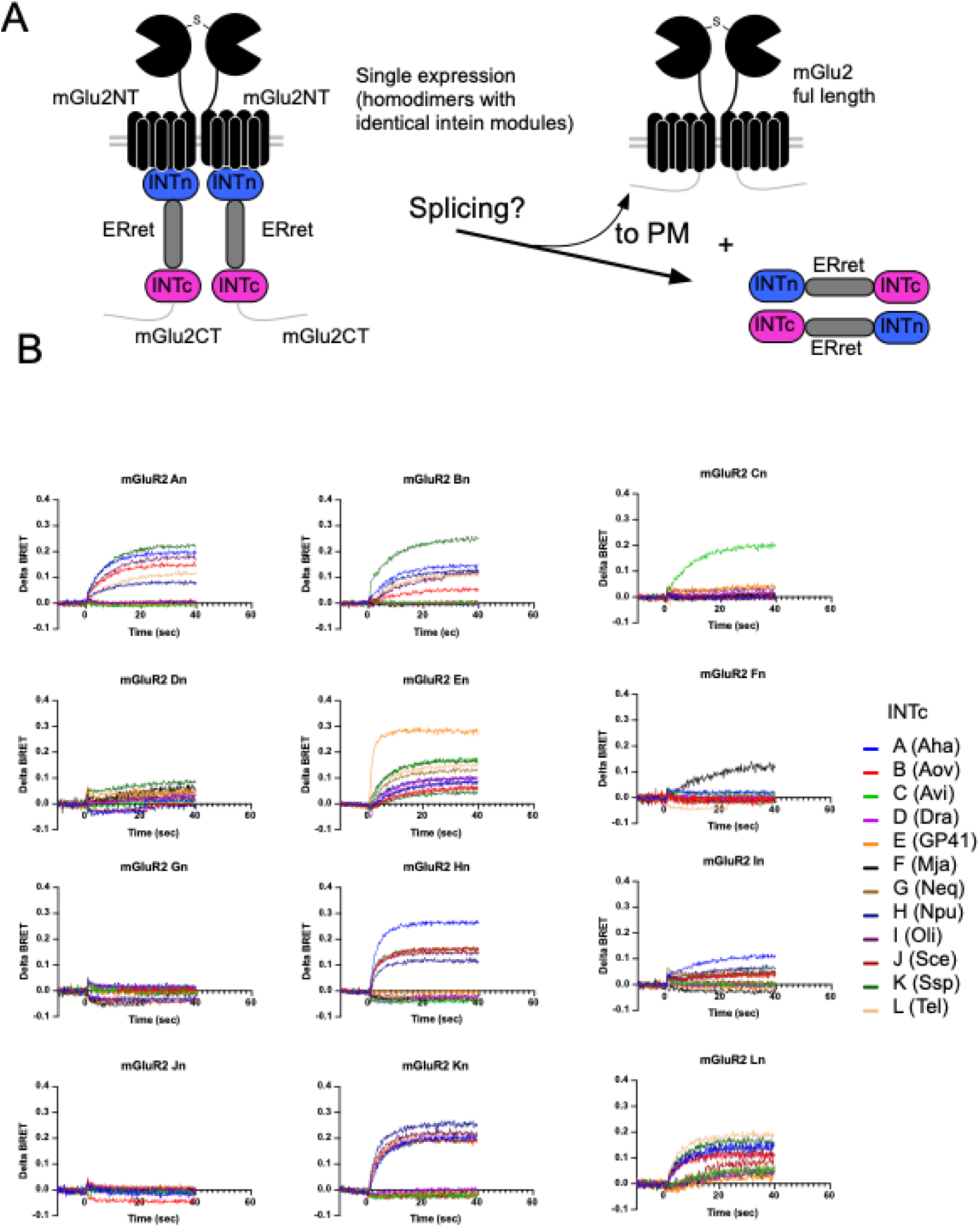
mGlu2 tagged with ER retention/internalization module flanked by 12 different N-and C-inteins. A, schematic of intein/retention module strategy. 144 mGlu2 constructs were tagged with each of 12 N-inteins and each of 12 C-inteins flanking a GABA_B_ receptor 1 (GBR1) C-tail style ER retention/internalization cassette. B, Each tagged receptor was expressed alone and function was assayed with the NanoBRET system (see text) by applying 1 mM glutamate at time = 0. Function was an indicator of surface expression and by extension, intein splicing. Thus in this assay, function was an indication that the given N-and C-intein pair were unsuitable for the assay moving forward. Note the key (right) indicating the single letter code indicating specific inteins to be used in the remaining figures.

The next step was to begin cloning the intein/ER retention modules into our receptors. As a starting point, we cloned every combination of the 12 N-and C-inteins flanking our retention sequence into the P1.1 position of mGlu2 (Fig. 2A) and expressed each receptor alone in HEK293T cells to be assayed using the NanoBRET system. For brevity, we created a shorthand nomenclature where each intein module was given a representative letter from A to L (see the key in Fig. 2B). Each letter indicates a specific intein and the n or c subscript (e.g., A_n_, or C_c_) denotes the N-or C-intein of that identity. Each panel in Fig. 2B shows the baseline and change in BRET upon application of 1 mM glutamate at time 0. When expressed alone, each receptor/intein module should only produce a change in BRET in response to glutamate if the retention sequence is spliced out and the mGlu2 homodimer is then trafficked to the plasma membrane. Detection of a change in BRET therefore indicates that splicing can occur between the N-and C-intein in the expressed receptor. Thus our expectation is that when an N-and C-intein in each receptor are compatible, we should see some signal, and that signaling in this assay should at least generally reflect the BRET signals seen in the soluble sensor screen described in Fig. 1. Indeed we found that that screen was a good predictor of the results obtained here. For example, only two inteins were ‘perfectly’ orthogonal: C and F (Avi and Mja from Fig. 1). In addition two inteins, D (Dra) and G (Neq) showed no evidence of splicing with any partner in either assay. With some inteins, namely E (GP41) and L (Tel), detectable BRET changes were evident with every other intein, although this was not the case in the soluble screen. This may suggest that the receptor signaling assay may be more sensitive than the soluble sensor assay, or that the ER retention system has some inherent ‘leak’ that is more pronounced with some intein modules more than others.

We next moved forward with several of the candidate mGlu2-intein modules that were non-responsive in this assay, indicating that these N-and C-intein pairs were mutually orthogonal, and could be usable in the subsequent studies. These were each expressed alone, and in pairs to determine if co-expression would result in splicing and plasma membrane expression, and consequently function (Fig. 3). Fig. 3A shows the results of one example experiment in which mGlu2CA (containing the module C_n_-ER retention-A_c_ plus the mGlu2 C-tail) and mGlu2AC were expressed alone, then both were expressed together. As predicted by the individual experiments in Fig. 2, neither mGlu2AC nor mGlu2CA exhibited any function upon 1 mM glutamate application, but when both were expressed a robust delta BRET was observed (Fig. 3A). Fig. 3B sows a summary of a number of other small scale experiments with other intein pairs expressed alone (red and blue) or together (green), suggesting that several pairs of inteins may be suitable for subsequent screening.

**Figure 3.**
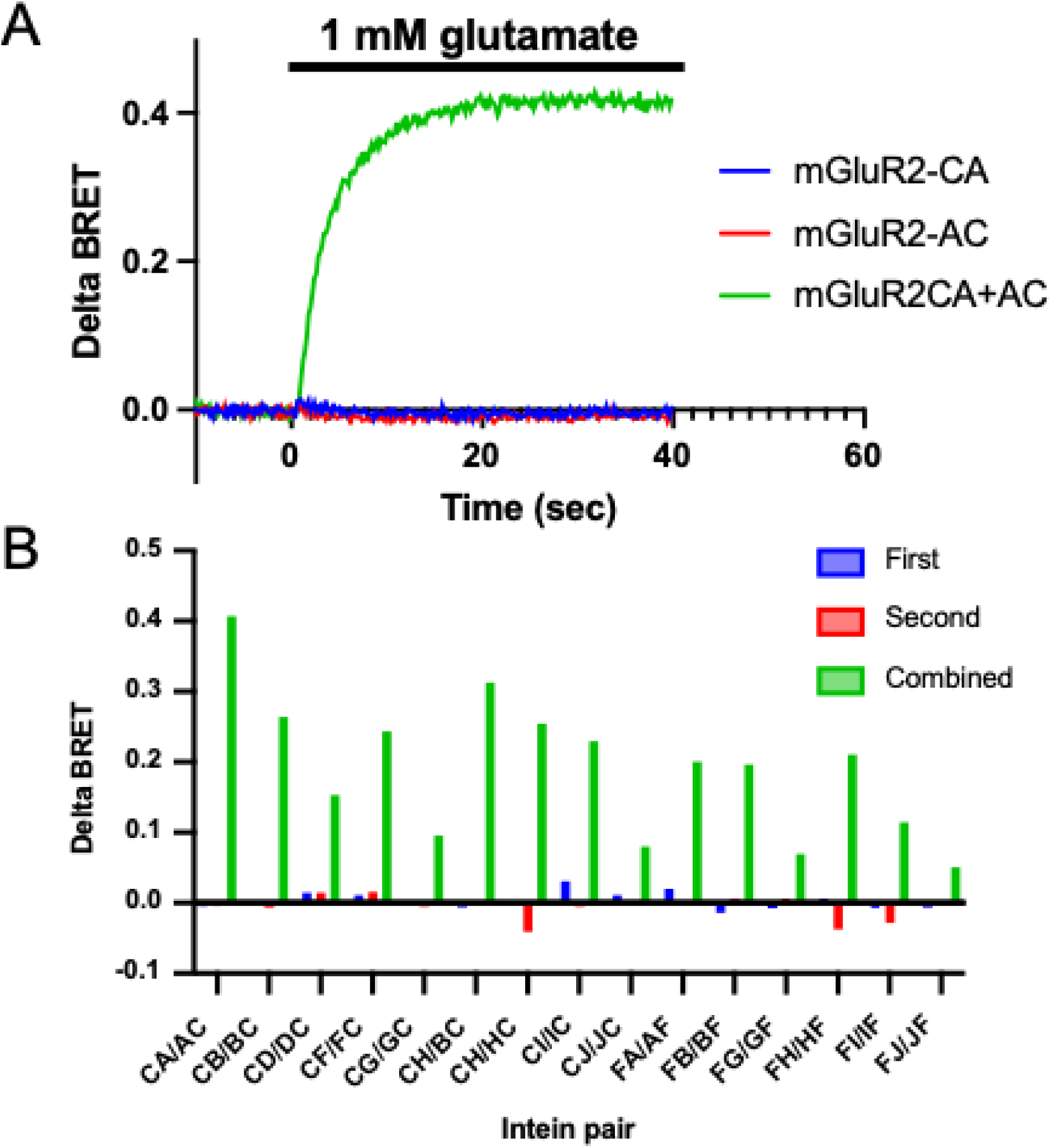
Responses of mGlu2 with paired intein/retention modules. A, Sample BRET signal induced by 1 mM glutamate in a HEK293T cell expressing mGlu2-CA-2ct (blue) or mGlu2-AC-2ct (red) constructs alone, or together (green). B, Average delta BRET from 2 experiments as in A, with mGlu2 tagged with the indicated intein/retention modules. C, Immunoblot showing the results of a biotinylation/pull-down experiment when full length SNAP-tagged mGlu2 is expressed, or when either mGlu2-EH-2ct or mGlu2-HE-2ct are expressed alone or together. I indicates input lane (cell lysates), S indicates supernatant, and IP indicates pull-down with avatin beads. Anti-GFP was used as a loading control (*below*). Arrows indicate the expected molecular weights of the unspliced and spliced SNAO-mGlu2 monomer bands, as shown. Note that a clear spliced band is only evident when the EH/HE pair is co-expressed, and that this band closely matches the molecular weight of the SNAP-mGlu2 construct with no C-terminal tags.

### Pharmacology of spliced dimers

Although several combinations of inteins within our retention module can efficiently prevent surface expression when expressed alone and promote it when combined, it is important to be cautious about possible false positive results. This could potentially occur if co-expression of receptors with compatible intein/retention modules results in close proximity of homodimers, leading to splicing and plasma membrane trafficking of one or both homodimer. This is difficult to assess when only a single receptor subtype is tagged, as in Fig. 3, because both the ‘homo’ and ‘hetero’ dimers will have identical pharmacology. Thus, we generated mGlu2 and 4 receptors tagged with several compatible intein/retention modules and examined selective orthosteric agonist responses to assess whether the surface receptors contained substantial numbers of homodimers. This receptor combination was chosen because mGlu2/4 dimers have known, unique pharmacology compared with each homodimer when exposed to the selective agonists DCG-IV (mGlu2) and L-AP4 (mGlu4), which act as very weak partial agonists, and either glutamate or the selective agonists applied together, which should act as full agonists^15,26^. In other words, strong activation by either selective agonist indicates that one or another receptor homodimer is reaching the plasma membrane. Fig. 4 shows the results of these experiments for 15 combinations of intein/retention module pairs. Although the CA/AC pair showed particularly strong responses to each selective agonist, the other combinations all showed profiles expected for a mainly heterodimer population: weak activation by each selective agonist, and all or near full agonist activity when combined. A few combinations, namely FG-GF and FJ-JF, showed weak or undetectable responses, suggesting that expression levels of the surface heterodimers may be low compared to that with other combinations.

**Figure 4.**
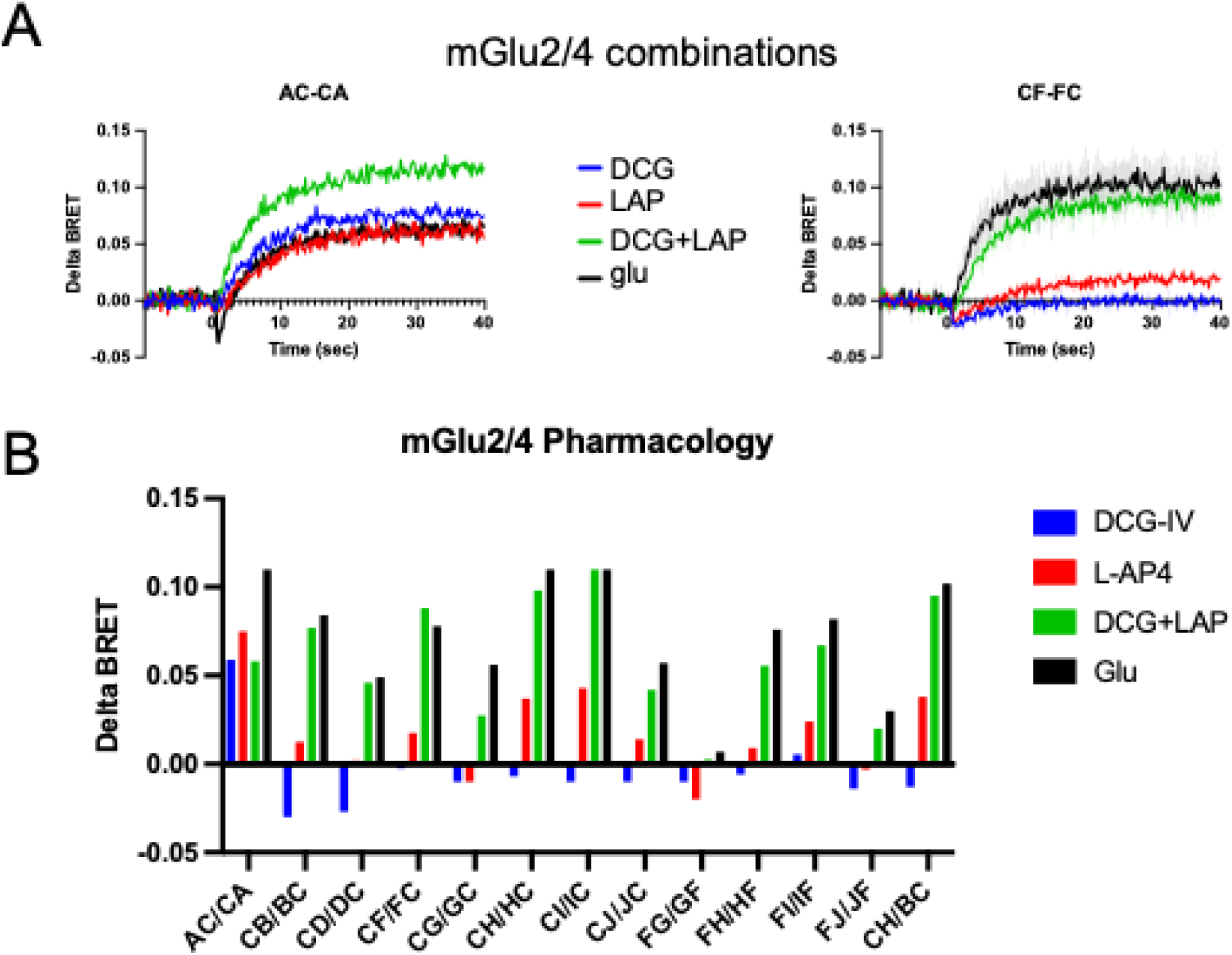
Several pairs of mGlu2/4 combinations with paired, compatible intein/retention modules (each mGlu2 has an mGlu4ct and each mGlu4 has an mGlu2ct) exposed to four agonist applications: 1 mM glutamate (black), 1 µM DCG-IV (blue), 30 µM L-AP4 (red), or DCG-IV + L-AP4 together (green). The label above each panel indicates the mGlu2 construct first and the mGLu4 construct second. For example, “AC-CA” indicates mGlu2-AC-4ct plus mGlu4-CA-2ct expressed together. Each value represents a single replicate.

### CF/FC intein/retention modules yield several heterodimer pairs

Because the C and F inteins showed very good orthogonality in the soluble BRET screen as well as in the receptor screen, we began by cloning these combinations into each of the mGluRs, expressed each with its appropriate pair (mG2CF3 + mG3FC2, mG2CF4 + mG4FC2… etc.), and generated glutamate concentration response curves using the NanoBRET assay ^21^. Receptors expressed individually did not produce detectable delat-BRET responses (not shown). Fig. 5 shows the results of these experiments. Fig. 5A shows the results when combining each mGlu2CFx with the corresponding mGluxFC2 construct, showing the full complement of mGlu2 containing dimers. Combining mGlu2CF1 and mGlu2CF5 with mGlu1FC2 and mGlu5FC2, respectively, did not yield any glutamate responses, as expected since these receptors are not predicted to form heterodimers ^8^. Each combination of mGlu2 with group II and III mGluRs did produce responses with somewhat different potencies and efficacies to glutamate. Similarly, full dimer panels were generated for each combination of mGlu3CFx, mGlu4CFx, mGlu7CFx, and mGlu8CFx, and these are shown in Figs. 5B-E. Many, but not all of the resulting curves strongly resemble those expected for the intended heterodimer pair ^15^. A few results were however, unexpected. For example, the expected group I mGluR pairings, mGlu1CFx and mGlu5CFx with the corresponding mGluxFC1 or 5 did not yield detectable delta BRET signals with the exception of mGlu5CF8+mGlu8FC5, including the homodimer pairings (1/1 and 5/5). It is unclear why pairings that included the group I receptors yielded poor results, but it may be due to some issues with their much larger intracellular C-tails compared with the group II and III receptors, or that they may require larger receptor numbers on the cell surface to yield measurable responses. We explore the group I receptors in more detail below. In addition, pairing mGlu8CF1 and mGlu8CF5 with mGlu1FC8 and mGlu5FC8 each resulted in measurable signals (Fig. 5E), despite these pairs being unlikely to heterodimerize ^8^.

**Figure 5.**
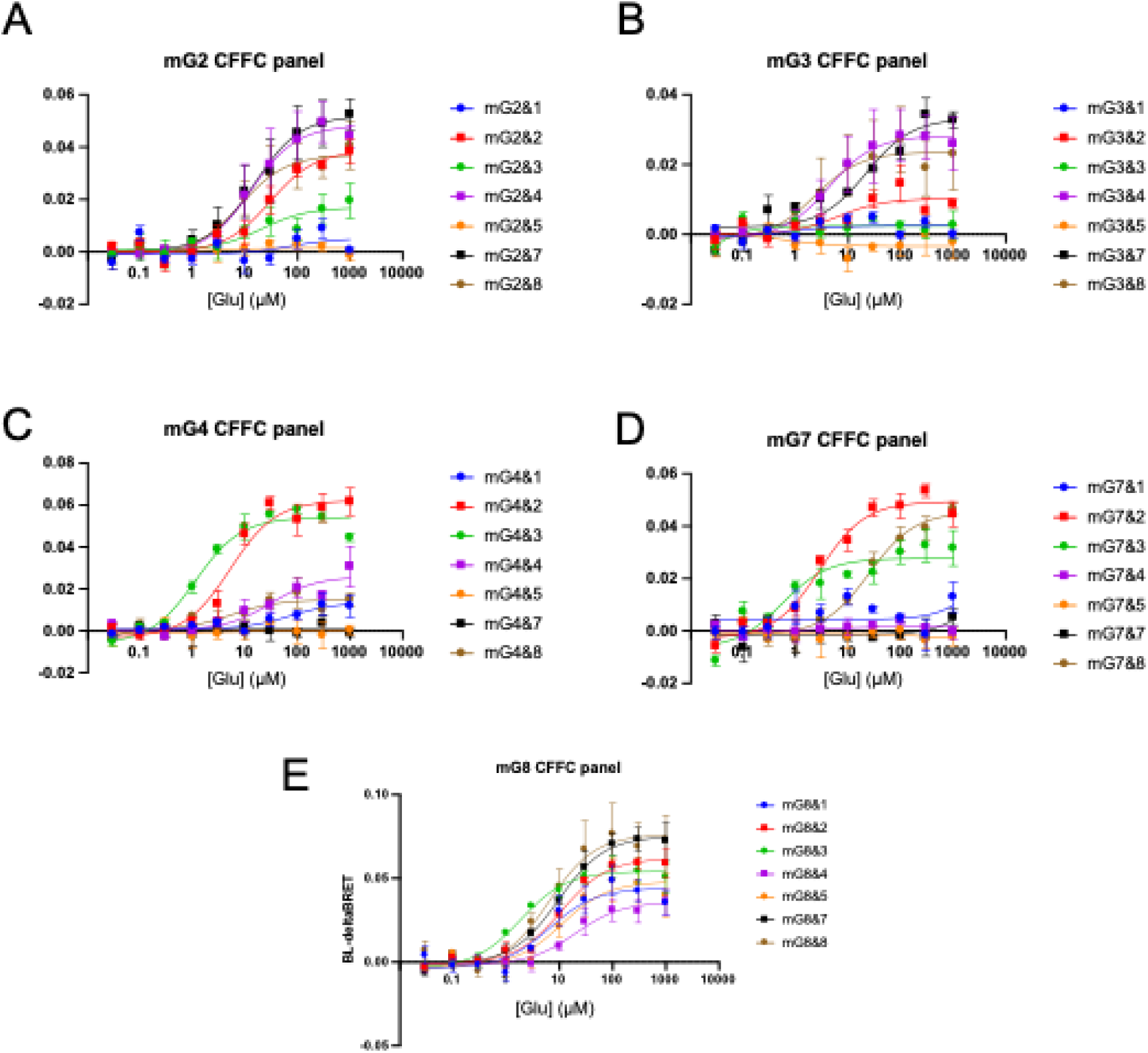
Glutamate dose-response curves for each combination of group II and III mGluRs with CF/FC intein modules. A, mGlu2-CF-x coexpressed with each mGlux-FC-2. Glutamate (30 nM-1 mM) was applied and ΔBRET is shown using the NanoBRET assay. Results from mGlu3-CF-x, mGlu4-CF-x, mGlu7-CF-x, and mGlu8-CF-x are shown in panels B-E, as indicated.

To directly examine the performance of the ER retention sequence used in the intein/retention modules, several receptors were tagged with an extracellular HiBit tag on the N-terminal end (Fig. 6A), and probed for both surface and total expression using purified LgBit protein. First, we applied purified LgBit protein^27^ to cells with the luciferase substrate Hikarazine to assess surface expression, then added 4 mg/ml digitonin to permeabilize cells, which allowed LgBit into the cells to assess total expression (Fig. 6B). As controls, similar measurements were made from untransfected cells, and cells expressing HiBit-tagged β-Arrestin, representing a pool of entirely intracellular HiBit tags. As a positive control, we will also express a full length HiBit-mGlu2. We made a panel of mGlu2-CF-x with N-terminal HiBit tags, and tested surface expression in this way (Fig. 6C). Application of the LgBit protein revealed that when expressed alone, several HiBit-mGlu2-CF-x receptors did exhibit some surface expression, although surface expression levels as a fraction of total were lower than HiBit-mGlu2 and increased significantly when the appropriate mGlux-FC-2 construct was co-expressed (with the exception of HiBit-mGlu2CF8). These data reveal that the retention/internalization motif was largely working, but suggested that some ‘leak’ was apparent. We also generated HiBit-mGlu7CF8 and HiBit-mGlu8CF7 (Fig. 6D). With these constructs, we noticed a dramatic difference in surface expression when expressed alone. While HiBit-mGlu7CF8 performed comparably to the HiBit-mGlu2 clones, HiBit-mGlu8CF7 showed strong surface expression when expressed alone, comparable to the HiBit-mGlu2 control (Fig. 6D). When its partner mGlu7FC8 was coexpressed, no additional surface expression was observed. These data suggest that the effectiveness of the retention sequence can depend on the backbone receptor it’s attached to, with mGlu8 receptors being inefficiently retained. This result can help explain the promiscuous signaling observed with all of the mGlu8CF clones described in Fig. 5 and suggest that it is likely mGlu8 homodimers may be escaping. Indeed, these constructs were the only ones that showed detectable signaling when expressed alone (Fig. 6D).

**Figure 6.**
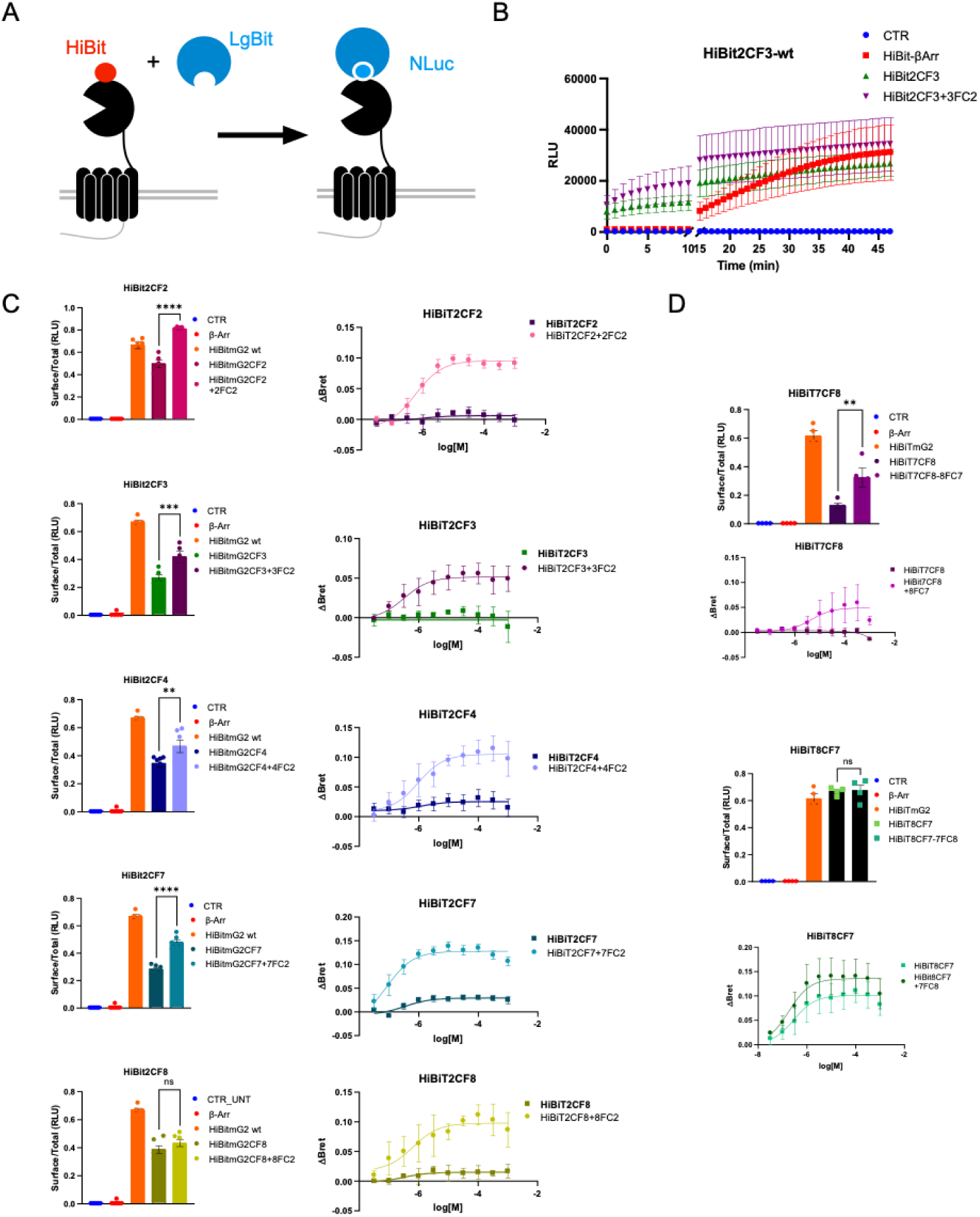
Assessment of ER retention and effects on signaling. A, Schematic showing the HiBit tag, LgBit surface expression strategy. Cells were exposed to LgBit protein and the luciferase substrate Hikarizine approximately 5 minutes before time =0. At 10 minutes, digitonin (4 mg/ml) was added to permeabilize cells to allow intracellular access to the LgBit protein so that total expression could be assessed. B, Time course of luminescence for surface and total HiBit expression. Negative control responses (untransfected KEK293t cells) are shown in blue. HiBit-ßArrestin served as the internal control (red). A shows results with the unmutated ER retention/internalization motif on the Hi-Bit tagged mGlu2CF3 construct as used in previous experiments. B shows the results from a similar experiment but with the retention motif mutated (“VA/ASVA”). C shows the ratio of surface to total expression for the indicated constructs for the entire mGlu2 panel (mGlu2CF paired with each mGluFC2 construct) with the unmutated retention motifs on the left and mutated on the right. Each experiments were run with negative control (untransfected cells in blue), an internal control (HiBit-ßArrestin cells in red), and a positive control (full length HiBit-mGlu2 with no intein module shown in orange). D shows responses to glutamate for each indicated expression condition.

### Assessing heterodimer pharmacology

To assess whether the receptors expressed with the group II and III mGluR CF/FC intein modules in fact led to a strict population of heterodimers, we compared the glutamate potencies of each resulting pair with the heterodimer specific G protein recruitment assay CODA-RET2, described in our recent paper ^15^. In the CODA-RET2 assay, mGluRs were tagged with either the large Bit (LgBit) or small bit (SmBit) fragments of Nanoluciferase which when assembled (with micromolar affinity) within an active receptor can lead to a BRET signal when a mini G protein tagged with Venus is co-expressed, thus giving a heterodimer specific BRET signal when the receptor is activated. Glutamate dose-response curves from that study (reproduced here with permission) ^15^, were normalized to peak responses and plotted against normalized responses obtained from co-expression of the group II and III mGluRs tagged with our intein/retention modules (Fig. 7). In most cases, expression of the selected CF/FC intein pairs produced responses whose glutamate potencies and slopes closely matched those obtained with the CODA-RET2 assay. The primary exceptions were the mGlu2/3 and mGlu4/8 combinations. In each case, this discrepancy is likely dut to the low efficacy of the responses using the intein approach, which led to a relatively noisy curve (Fig. 7A and H). A few receptor combinations were also examined with the AC/CA inteins, but these yielded less satisfactory results than the CF/FC combination (Fig. 7 G-I).

**Figure 7.**
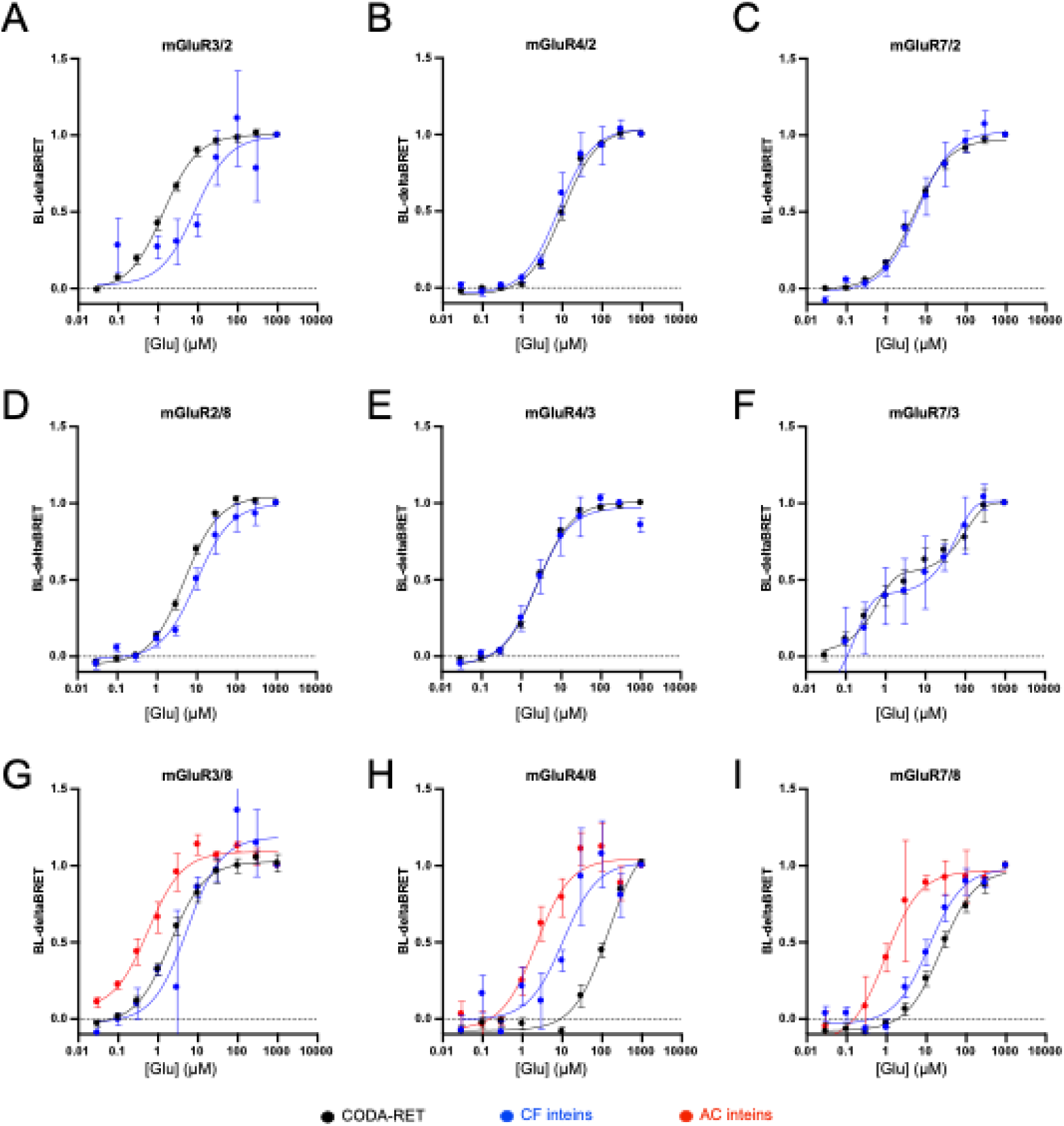
Comparison of glutamate responses using the intein/retention strategy (from Fig. 5) to those obtained using the CODA-RET2 assay, normalized to the peak responses. CODA-RET2 responses are shown in black, CFFC intein responses are shown in blue, and some ACCA intein responses are shown in red. The normalized CFFC intein responses are calculated from the combined reciprocal expression of the two CFFC pairs. For example, “mGlu3/2” included responses from both mGlu2-CF-3 + mGlu3-FC-2 and mGlu3-CF-2 + mGlu2-FC-3. Panels A-I show mGlu3/2, mGlu4/2, mGlu7/2, mGlu2/8, mGlu4/3, mGlu7/3, mGlu3/8, mGlu4/8, and mGlu7/8, respectively.

## Discussion

Pharmacological responses of mGluR heterodimers to selective ligands are difficult to predict from the pharmacologies of their corresponding homodimers^15^. For that reason, the effect of any novel orthosteric or allosteric ligand acting on an mGluR subunit should be tested on each possible heterodimer expressed in isolation. However, the tendency of each mGluR to form homodimers as well as heterodimers confounds this process. This is of particular importance as several mGluR targeting compounds have recently entered clinical trials without empirical data on how they affect the function of several of the heterodimeric receptors they potentially modulate^4,28,29^. Here we describe a new method to direct the selective expression of specific mGluR heterodimers using a strategy employing orthogonal, split inteins flanking a large ER retention/internalization motif in the proximal C-terminal tails of group II and III mGluRs. While this method is used here for selective expression of mGluR dimers, it may also be applicable to other dimeric cell surface receptors or even tetrameric receptors that form dimer-dimer tetramers such as NMDA receptors.

After an initial screen of several split inteins for orthogonality and cloning each into the N-and C-positions in our intein/retention module on the C-tail position of mGlu2, a smaller number were chosen for functional assessment and pharmacological screening when paired with mGlu4 to compare with known mGlu2/4 heterodimer pharmacology. From these, one intein pair, “CF” (AviPRP8 and MjaKlbA) were chosen for the full screen in which every mGluR excluding mGlu6 was tagged with intein modules containing the Cn and Fc inteins and Fn and Cc inteins flanking the retention sequences surrounded by each mGluR N-terminal domain and each C-tail. Each of these was then co-expressed such that the appropriate C-tail would be spliced on to each mGluR N-terminal domain and glutamate dose-response curves were generated from the resulting receptors. In most cases we observed considerable differences in the glutamate potencies consistent with signaling of the intended heterodimers (Fig. 7). Close examination of responses from each heterodimer pair reveals that mGlu2/3 and mGlu4/8 were the main combinations that did not respond as expected based on predictions from the CODA-RET2 approach^15^. The main issue with the mGlu2/3 combination is low efficacy (see Fig. 5), possibly due to low expression levels. This could be a consequence of low expression or potentially that the small ‘scar’ left after intein splicing is uniquely detrimental to the group II receptors. Indeed, the efficacies of not only mGlu2/3 (Fig. 5 ACB) but also mGlu2/2 (Fig. 5A) and mGlu3/3 (Fig. 5B) were all lower than expected. Thus, we cannot say with certainty whether the underlying basis of the observed low efficacy of the group II combinations is scar location or just that intein splicing is inefficient with these combinations. This could necessitate further optimization in the future.

With respect to the mGlu4/8 dimers, the resulting response appears left shifted compared with the expected response. This is likely due to ‘leak’ of mGlu8 homodimers, which is likely due to inefficient ER retention when the mGlu8 N-terminal domain precedes the CF module, as shown in Fig. 6. Fig. S3A shows a comparison of each configuration of these results (in Fig. 7 both combinations are averaged together), using an arbitrarily scaled CODA-RET trace against the BRET values obtained with both mGlu4FC8+8FC4 and mGlu8CF4+4FC8. These data suggest that neither combination strongly reflects the heterodimer response predicted with the CODA-RET method.

While it isn’t clear why retention might differ with mGlu8CF constructs vs. mGlu8FC constructs, in the case of the 7/8 heterodimers constructed in each way, responses with mGlu7CF8+8FC7 seem to reflect the CODA-RET results better than mGlu8CF7+7FC8 (see Fig. S3B, which compares against a scaled CODA-RET response, keeping in mind that the Emax value here is not meaningful across methods). Thus, it may be generally favorable to rely more on combinations using mGlu8FC rather than mGlu8CF constructs, which may be retained better, or just lead to more efficient intein splicing. Together these results highlight the context dependence of not only retention efficiency but also possibly intein splicing. Lastly, we note that with the mGlu4/7 combination we did not observe any discernable signaling with glutamate concentrations up to 1 mM. However, even with the CODA-RET approach discussed above^15^, this also combination failed to yield meaningful signaling in this range (not shown).

Here we have presented a novel strategy for achieving composition controlled selective heterodimer expression using a combination of ER retention/internalization sequences and orthogonal, split inteins. We have found that for most combinations of group II and III mGluRs, this system is able to produce functional, isolated heterodimer expression that reflects predicted heterodimer pharmacology. With further optimization, it is likely that pure populations of the resistant heterodimers such as mGlu4/8 could also be achieved. Finally, this strategy should be adaptable to other receptors or membrane proteins in which homo-and heterodimers, or dimer-dimer tetramers are necessary for function.

## Supporting information

Supplemental MaterialsFig

