## Supplemental MaterialsFig for "Use of orthogonal split inteins to direct surface expression of specific metabotropic glutamate receptor hererodimer pairs"

### Supplemental Materials

Figure S1. Titration and rapamycin effects of soluble intein constructs. A, *upper* shows western blot labeling the N-intein fragment (anti-HA in green) and the C-intein fragment (anti-NLuc in red), as in Figure 1, as the amount of cDNAs were varied as shown . Panels in the *center* show each blot separately. *Lower* panel shows BRET signal from each transfection shows that the BRET value correlated with the splice product band labeled with both antibodies (arrow). B, Only two intein constucts showed rapamycin dependent BRET signals. The Neq$_C$ intein (*upper*) exhibited enhanced BRET with most N-inteins, although no splice product was observed for any. Sce$_C$ also exhibited rapamycin dependent BRET signals with some N-inteins.

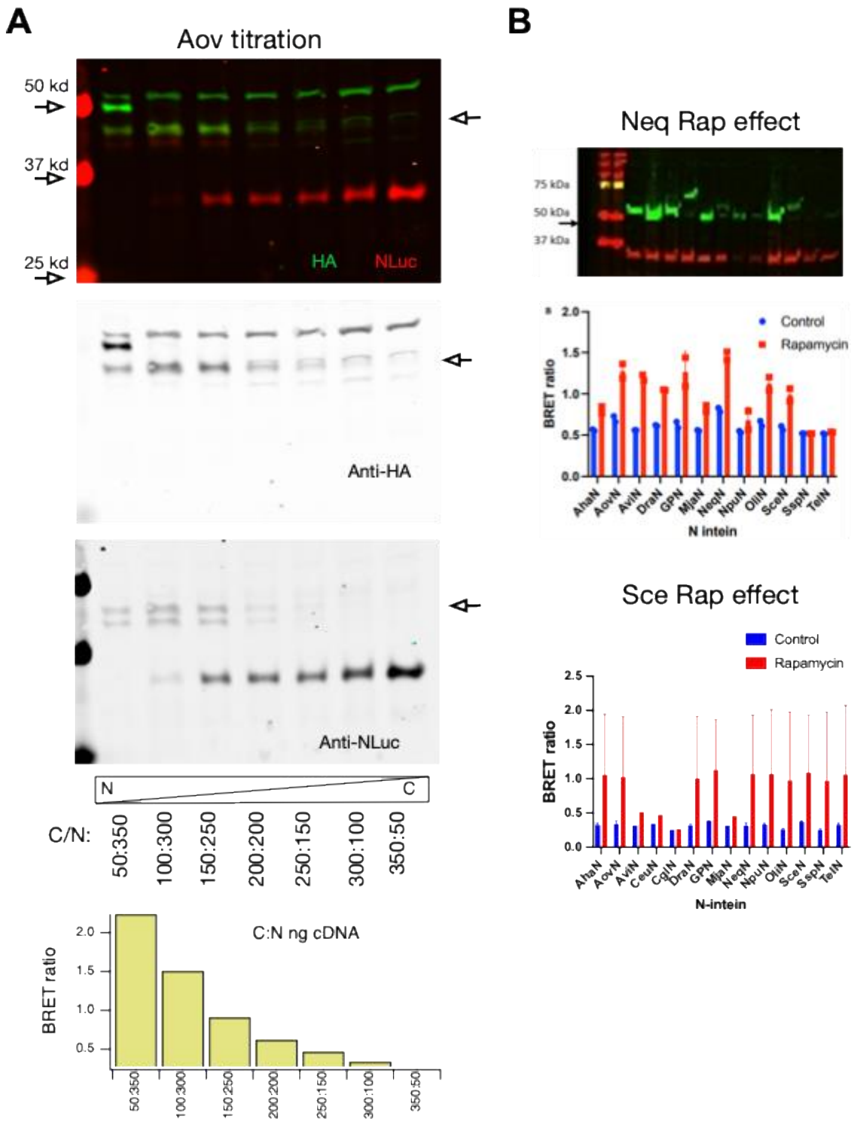

Figure S2. Effect of intein scar position. The residual ‘extra’ amino acids (AEY|CFN) predicted to be left after Npu intein splicing were cloned into 5 positions in the mGlu1 C-tail, labeled “Pos 1.0” through 1.4 (A and B). Position 1.0 was immediately after the 7$^{th}$ transmembrane domain while each subsequent position pas +5 amino acids. Panel A shows that position 1.0 seemed to occlude mGlu1 function while each of the other positions was permissive for function (A and B). Panel C shows the effect of different scars (as indicated) at position 1.1 on the glutamate dose-response curve of mGlu1. Scars used were Npu, Avi (FWERA|GFEE), and Mja (TGHDG|SGTL).

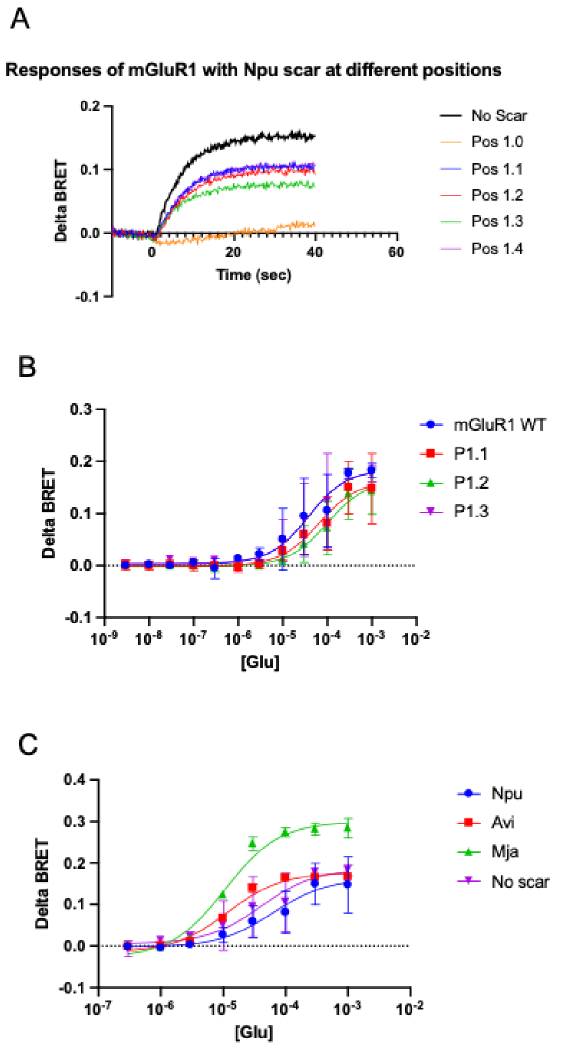

Figure S3. Positional effects of C&F inteins in two heterodimer pairs. A, shows glutamate dose-response curves resulting from the CODA-RET method (*black*, arbitrarily normalized) and by combining mGlu4CF8 with mGlu8FC4 (green) or mGlu8CF4 with mGlu4FC8 (magenta). B, shows glutamate dose-response curves resulting from the CODA-RET method (*black*, arbitrarily normalized) and by combining mGlu7CF8 with mGlu8FC7 (green) or mGlu8CF7 with mGlu7FC8 (magenta).

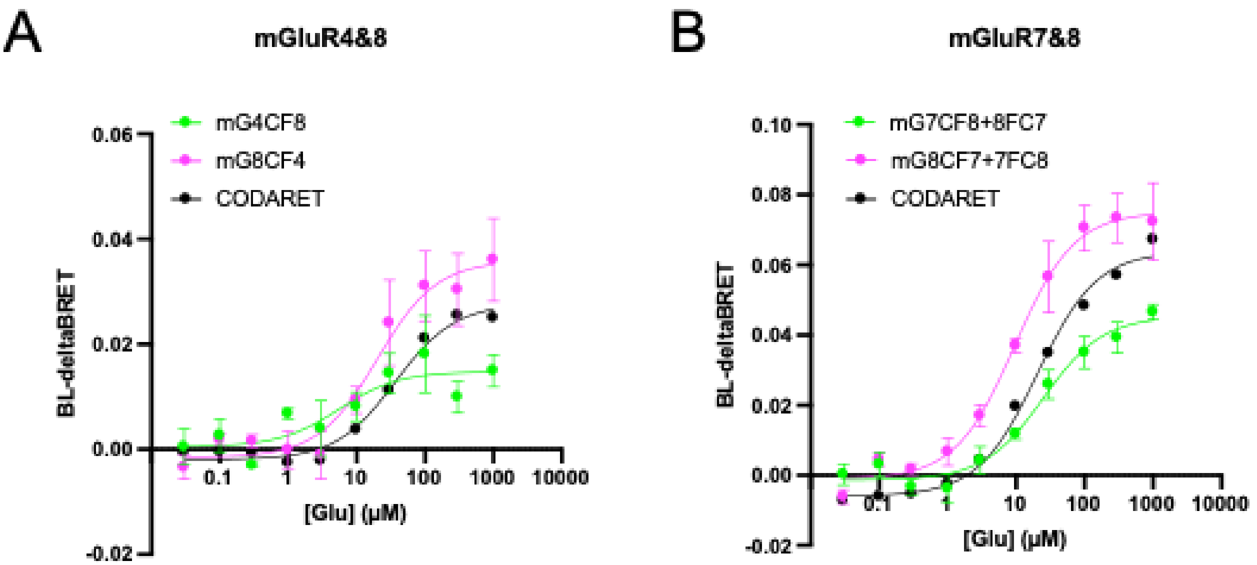

Appendix 1. Amino acid sequences of intein and ER retention/internalization motifs.

| N-Intein construct | Symbol | AA Sequence |
|---|---|---|
| Aha DnaE | $A_n$ | AEYCLSYDTEIWTVEYGAMPIGKIVEEKIECSVYTVDENGFVYTQPIAQWHPRGQQEIIEYTLEDGRKIRATKDHKMMTESGEMLPI EEIFQRELDLKVETFHEMSLLRRGAK |
| Aov DnaE | $B_n$ | AEYCLSADTEILTVEYGFLPIGEIVGKAIECRVYSVDGNGNIYTQSIAQWHNRGEQEVFEYTLEDGSIIRATKDHKFMTTDGEMLPID EIFARQLDLMQVQGLH |
| Avi PRP8 | $C_n$ | FWERACLAKGTRLLRYDGSEIEVQDVKEGDLLLGPDGGPRRVFNIVNGEDRLYRIKIDRGIEDLVVTPNHILVLHREKKAKNNEDD ELPEVPAAERYD |
| Dra Snf2 | $D_n$ | MGLGKAQPLDAKVLTPLGWRTMGELQVGDYVIGRNGQPTQIVGVYPQGERPIYRVTLTDGASVEADAEHLWNVNTPVRKKRGLP ERTLTTAEIAADLQDAAGNLKHYLPLVEPVQFAGRDLPLDPYTLGALLGDGCFVHGVEITSEDELVSALPLPAGVEARLGTRLTPTV STSRLVTAGQWTPNPLKDALRGLGLHGKSGRYKFIPPDYLLGSPAQPAGSAARTPRH |
| GP41-1 | $E_n$ | TRSGYCLDLKTQVQTPQGMKEISNIQVGDLVLSNTGYNEVLNVFPKSKKKSYKITLEDGKEIICSEEHLFPTQTGEMNISGGLKEG MCLYVKEI |
| Mja KlbA | $F_n$ | TGHDGALAYDEPIYLSDGNIINIGEFVDKFFKKYKNSIKKEDNGFGWIDIGNENIYIKSFNKLSLIIEDKRILRVWRKKYSGKLIKITTK NRREITLTHDHPVYISKTGEVLEINAE |
| Neq Pol | $G_n$ | VIYGDSIMDTEIEVIENGIKKKEKLSDLFNKYYAGFQIGEKHYAFPPDLYVYDGERWVKVYSIIKHETETDLYEINGITLSANHLVLSK GNWVKAKEYENKNN |
| Npu GyrB | $H_n$ | AEYCLSYETEILTVEYGLLPIGKIVEKRIECTVYSVDNNGNIYTQPVAQWHDRGEQEVFEYCLEDGSLIRATKDHKFMTVDGQMLPI DEIFERELDLMRVDNLPN |
| Oli DnaE | $I_n$ | AEYCLSYNTEVLTVEYGPLPIGKIVDEQIHCRVYSVDENGFVYTQAIAQWHDRGYQEIFAYELADGSVIRATKDHQFMTEDGQMFP IDEIWEKGLDLKKLPTVQDLPAAVGYTVS |
| Sce VMA | $J_n$ | IIYVGCFAKGTNVLMADGSIECIENIEVGNKVMGKDGRPREVIKLPRGRETMYSVVQKSQHRAHKSDSSREVPELLKFTCNATHE LVVRTPRSVRRLSRTIKGVEYFEVITFEMGQKKAPDGRIVELVKEVSKSYPISEGPERANELVESYRKASNKAYFEWTIEARDLSLLG SHVRKATYQTYAPILY |
| Ssp GyrB | $K_n$ | AEYCLSFGTEILTVEYGPLPIGKIVSEEINCSVYSVDPEGRVYTQAIAQWHDRGEQEVLEYELEDGSVIRATSDHRFLTTDYQLLAIE EIFARQLDLLTLENIKQTEEALDNHRLPFPLLDAGTIK |
| Tel DnaE | $L_n$ | AEYCLSGETAVMTVEYGAVPIRRLVQERLSCHVYSLDGQGHLYTQPIAQWHFQGFRPVYEYQLEDGSTICATPDHRFMTTRGQML PIEQIFQEGLELWQVAIAPRQALLQGLKPAVQMSG |
| **Retention/Internalization motif (GBR CT)** | | gsggTMKTGSSTNNNEEEKSR**LL**EKENRELEKIIAEKEERVSELRHQLQSRQQL**RSRR**HPPTPPDPSGGLPRGPSEPPDRLSCDG SRVHLLYK |
| **C-Intein construct** | **Symbol** | **AA Sequence** |
| Aha DnaE | $A_c$ | MVKIIKRQSLGRQNVYDVCVETDHNFVLANGCVASNCFN |
| Aov DnaE | $B_c$ | MVKITARKFVGRENVYDIGVEHHHNFAIKNGLIASNCFN |
| Avi PRP8 | $C_c$ | TVEMTAAEFAALSTEERSLYRAFRCPGFELSEKAVPANHSFTIKDIRLELETTEWAGFRVDKDQLYLRHDYLVLHNSGFEE |
| Dra Snf2 | $D_c$ | MGWGCMARAGATNSFPRITCWEAPRSRLAVLQGLLDTDGHAGVVVEYVSVSEHLARGVVELVQSLGGVARIRQKATSHTYGGE KKTGLAWRATLKLPPELEPFRLAAKRAAYRRPTKYPPTRGIKSIEWVGHKPAQCIAVDAPDHLYVTEGYIVTHNTLQTL |
| GP41-1 | $E_c$ | MMLKKILKIEELDERELIDIEVSGNHLFYANDILTHNSSSDV |
| Mja KlbA | $F_c$ | MVKVGDYIYIPKNNTINLDEVIKVETVDYNGHIYDLTVEDNHTYIAGKNEGFAVSNCSGTL |
| Neq Pol | $G_c$ | MRYLGKKRVILYDLSTESGKFYVNGLVLHNTDSLF |
| Npu GyrB | $H_c$ | IKIATRKYLGKQNVYDIGVERDHNFALKNGFIASNCFN |
| Oli DnaE | $I_c$ | MVKIVRRQSLGVQNVYDIGVEKDHNFCLASGEIASNCFN |
| Sce VMA | $J_c$ | VLLNVLSKCAGSKKFRPAPAAAFARECRGFYFELQELKEDDYYGITLSDDSDHQFLLANQVVVHN |
| Ssp GyrB | $K_c$ | MVKVIGRRSLGVQRIFDIGLPQDHNFLLANGAIAANCFN |
| Tel DnaE | $L_c$ | MKIVGRRLMGWQAVYDIGLAADHNFVLANGAIAANCFN |
